# Hybrid origins and the earliest stages of diploidization in the highly successful recent polyploid *Capsella bursa-pastoris*

**DOI:** 10.1101/006783

**Authors:** Gavin M. Douglas, Gesseca Gos, Kim A. Steige, Adriana Salcedo, Karl Holm, Emily B. Josephs, Ramesh Arunkumar, J. Arvid Ågren, Khaled M. Hazzouri, Wei Wang, Adrian E. Platts, Robert J. Williamson, Barbara Neuffer, Martin Lascoux, Tanja Slotte, Stephen I. Wright

**Affiliations:** Department of Ecology and Evolutionary Biology, University of Toronto, Toronto, Canada; Department of Ecology and Genetics, Evolutionary Biology Centre, Science for life Laboratory, Uppsala University, Uppsala, Sweden; Center for Genomics and Systems Biology, New York University Abu Dhabi, Abu Dhabi, United Arab Emirates; McGill Centre for Bioinformatics, McGill University, Montreal, Canada; Department of Botany, University of Osnabrück, Osnabrück, Germany; Department of Ecology, Environment and Plant Sciences, Science for Life Laboratory, Stockholm University, Stockholm, Sweden

**Keywords:** polyploidy, population genomics, speciation, gene loss

## Abstract

Whole genome duplication events have occurred repeatedly during flowering plant evolution, and there is growing evidence for predictable patterns of gene retention and loss following polyploidization. Despite these important insights, the rate and processes governing the earliest stages of diploidization remain poorly understood, and the relative importance of genetic drift, positive selection and relaxed purifying selection in the process of gene degeneration and loss is unclear. Here, we conduct whole genome resequencing in *Capsella bursa-pastoris*, a recently formed tetraploid with one of the most widespread species distributions of any angiosperm. Whole genome data provide strong support for recent hybrid origins of the tetraploid species within the last 100-300,000 years from two diploid progenitors in the *Capsella* genus. Major-effect inactivating mutations are frequent, but many were inherited from the parental species and show no evidence of being fixed by positive selection. Despite a lack of large-scale gene loss, we observe a decrease in the efficacy of natural selection genome-wide, due to the combined effects of demography, selfing and genome redundancy from whole genome duplication. Our results suggest that the earliest stages of diploidization are associated with quantitative genome-wide decreases in the strength and efficacy of selection rather than rapid gene loss, and that non-functionalization can receive a 'head start' through a legacy of deleterious variants and differential expression originating in parental diploid populations.

## SIGNIFICANCE

Plants have undergone repeated rounds of whole genome duplication, followed by gene degeneration and loss. Using whole genome resequencing we examined the origins of the recent tetraploid *Capsella bursa-pastoris* and the earliest stages of genome evolution after polyploidization. We conclude the species had a hybrid origin from two distinct *Capsella* lineages within the last 100-300,000 years. Our analyses suggest the absence of rapid gene loss, but provide evidence that the species has large numbers of inactivating mutations, many of which were inherited from the parental species. Our results suggest that genome evolution following polyploidy is determined not only by genome redundancy, but also by demography, mating system and the evolutionary history of the parental species.

## INTRODUCTION

Following whole genome duplication (WGD) events, genomes undergo a process of diploidization, where duplicate genes are lost, and only a minority of genes are retained as duplicates for extended periods of evolutionary time (1, 2). While this process has been well characterized (3), the dominant evolutionary forces driving gene degeneration and loss, and the rate at which diploidization occurs, remain unclear.

Large-scale genome projects have provided a number of important insights into genome evolution in ancient polyploids (1, 4, 5). First, gene retention and loss is nonrandom with respect to gene function, with signs of strong convergence in which genes are lost (1). This raises an open question about the extent to which gene loss proceeds solely via a process of relaxed purifying selection, or whether there is also positive selection for gene loss. Second, gene loss is often biased towards particular chromosomes, such that genes from one chromosome duplicated by WGD (homeolog) are preferentially lost (4), or become expressed at lower levels than those on the other homeolog (5). This process is termed biased fractionation (6), and while the underlying mechanism remains unresolved, it could result from differences in epigenetic regulation or selective history of the parental species (7), and may occur rapidly upon polyploid formation or following a longer timescale of genome rearrangement and evolution.

Studying the early stages of gene degeneration and loss should provide important insights into the causes of genome evolution in polyploids (8, 9) for several reasons. First, transitions to polyploidy are often accompanied by strong demographic bottlenecks and shifts to higher selfing rates (10), which can limit the efficacy of natural selection regardless of ploidy. Examination of population genomic patterns in recent polyploids can disentangle the impact of demography and gene redundancy on the efficacy of selection. Second, genomic studies of recent polyploids also allow inference about the extent of early gene degeneration following whole genome duplication. Rapid genome rearrangements and deletions have previously been identified in certain neopolyploids within even a single generation (11), and past work examining a small set of genes indicated that gene loss may be extremely rapid (9). Rapid divergence in expression between homeologous genes (“transcriptomic shock”) has also been identified following polyploidy (12, 13), although the relationship between transcriptomic shock and permanent genomic changes remains to be resolved. Overall, these studies imply that derived changes occur in the earliest stages of ploidy transitions; however the genome-wide extent of such changes remains unclear. Finally, whether the fate of genes in polyploid lineages is influenced by the evolutionary history prior to WGD, or solely through *de novo* changes following polyploidization remains a key open question (14). Comparative population genomics and expression analyses of early polyploids and their recent progenitor species provides an outstanding opportunity to better understand the dynamics associated with the early stages of polyploid evolution.

The genus *Capsella* includes the primarily self-fertilizing tetraploid *Capsella bursa-pastoris* (2n=4x=32) as well as three diploid species: the self-incompatible outcrosser *Capsella grandiflora*, and two self-compatible species *Capsella rubella* and *Capsella orientalis (Cg, Cr, Co:* 2n=2x=16). These species differ greatly in their geographical distribution. The outcrosser *C. grandiflora* is limited to north-western Greece and Albania, whereas the self-compatible *C. rubella* has a broader Mediterranean-central European distribution and self-compatible *C. orientalis* is found from eastern Europe to central Asia (15). In contrast to its diploid congeners, the selfing tetraploid *C. bursa-pastoris* has a worldwide distribution that is at least in part anthropogenic (16, 17).

We have previously shown that the self-fertilizing diploid *C. rubella* is derived from the outcrossing, self-incompatible diploid *C. grandiflora*, and that these species split relatively recently, within the last 50,000-100,000 years (18–21). There is reason to believe that outcrossing is the ancestral state in this system, as *C. grandiflora* shows trans-specific shared polymorphism with *Arabidopsis lyrata* and *Arabidopsis halleri* at the self-incompatibility locus (19, 22, 23). The self-compatible *C. orientalis* is therefore also thought to be derived from an outcrossing, *C. grandiflora-like* ancestor, although further back in time than *C. rubella* (15).

The origin of the widespread tetraploid *C. bursa-pastoris* has however proven difficult to determine (17, 24). Various hypotheses on the origin of this species have been put forward (15–16, 24–27), and most recently, *C. bursa-pastoris* has been suggested to be an autopolyploid of *C. orientalis* (15), or *C. grandiflora* (16). Despite the hypotheses of autopolyploidy, *C. bursa-pastoris* appears to have strict disomic inheritance, with two separate homeologous genomes segregating independently (26).

In this study, we investigate the mode of formation and genomic consequences of polyploidization in *C. bursa-pastoris* using high-coverage massively parallel genomic sequence data. We first evaluate the origins of *C. bursa-pastoris* and conclude that the species has recent, hybrid, allopolyploid origins from the *C. orientalis* and the *C. grandiflora/C. rubella* lineages.

By comparing genome-wide patterns of nucleotide polymorphism and gene inactivation between *C. bursa-pastoris* and its two close relatives, *C. grandiflora* and *C. orientalis*, we then quantify the rate and sources of early gene inactivation following polyploid formation. Our results suggest that the earliest stages of degeneration and gene silencing have been strongly influenced by selection and expression patterns in the diploid parents, highlighting a strong ‘parental legacy’ (14) in early polyploid genome evolution. Furthermore, relaxed selection in this early polyploid is driven not only by genomic redundancy due to ploidy, but also in large part by demographic effects and the transition to selfing. The results are important for an improved understanding of the consequences of polyploidization for the strength and direction of selection in plant genomes.

## RESULTS and DISCUSSION

### De Novo *Assemblies and Whole Genome Alignment Across the* Capsella *Genus*

We generated *de novo* assemblies for *C. bursa-pastoris* and *C. orientalis* (28) using Illumina genomic sequence (see Methods and SI Text). Scaffolds from these assemblies were then aligned to the *C. rubella* reference genome (21) using whole genome alignment, along with assemblies of *C. grandiflora* and the outgroup species *Neslia paniculata* (21). With the exception of *C. bursa-pastoris*, only a single orthologous chain was retained for each genomic region, while for *C. bursa-pastoris* we allowed up to two chains, based on the expectation of assembling sequences from the two homeologous chromosomes. In total, our *C. bursa-pastoris* assembly spans approximately 102MBase of the 130MBase *C. rubella* assembly, of which approximately 40% (42MBase) is covered by two homeologous sequences, while the remainder is covered by a single sequence.

While it is possible that regions covered by a single *C. bursa-pastoris* sequence reflect large-scale gene deletions, it is more likely that our assembly method often collapses the two homeologous sequences into a single assembled sequence, given the relatively low pairwise differences between the subgenomes (average of 3.7% differences in assembled subgenomes). For investigating the genomic relationships across the *Capsella* genus, we therefore focus on the 42MBase of the genome that contains two *C. bursa-pastoris* subgenomes, and provide further support that these regions are representative of genome-wide patterns in follow-up polymorphism analyses below.

### The Evolutionary Origin of Capsella bursa-pastoris

We constructed independent phylogenetic trees for each alignment from genomic fragments that included two homeologous regions of *C. bursa-pastoris,* and one each from *C. grandiflora*, *C. orientalis*, *C. rubella* and the outgroup sequence *N. paniculata*. We obtained 17,258 trees where all branches had at least 80% bootstrap support. The total alignment length across these trees represents 13.6 MBase of sequence across the genome. Of these trees, 70% showed clustering of one subgenome with the *C. grandiflora/C. rubella* lineage (Fig. 1 *A-C*; hereafter the ‘A’ subgenome) and the second subgenome with *C. orientalis* (hereafter the ‘B’ subgenome), providing strong support for a hybrid origin of *C. bursa-pastoris.* Thus, we conclude that *C. bursa-pastoris* is an allopolyploid resulting from a hybridization event between these two lineages, and these conclusions have also been supported by Sanger sequencing of a set of genes (SI text; Fig. S1). As *C. bursa-pastoris* clusters with *C. orientalis* for maternally inherited chloroplast DNA (15), we conclude that the maternal parent of *C. bursa-pastoris* came from the *C. orientalis* lineage and the paternal contribution came from an ancestral population from the *C. grandiflora*/*C. rubella* lineage. Given the current disjunct distribution of *C. grandiflora* and *C. orientalis* (15), this implies that the ancestral ranges must have been overlapping in the past, to allow for hybridization to occur.

**Figure 1.**
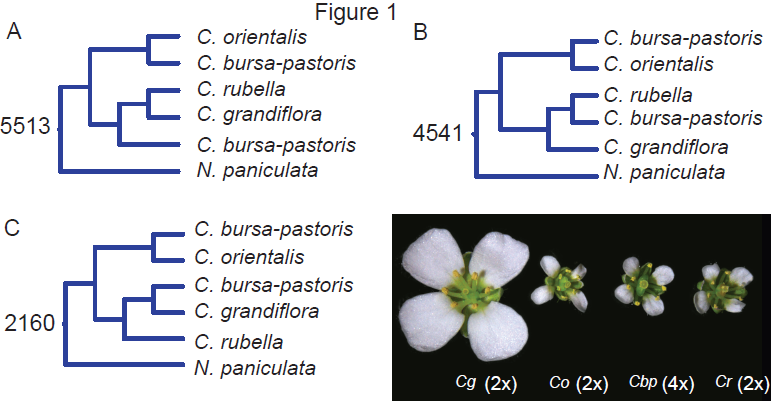
Results of RAxML tree analysis from whole genome alignments of the four *Capsella* species, using *Neslia paniculata* as an outgroup. Numbers show the number of trees with greater than 80% bootstrap support showing each most common topology (of 17,258 total), from genome-wide alignments.

Overall, our comparative 4-species genome analysis provides strong support for an allopolyploid origin of *C. bursa-pastoris.* To verify that these conclusions are not biased by a focus on genomic regions that have two subgenomes assembled, and to investigate polyploid origins in more detail, we resequenced 9 additional genomes of *C. bursa-pastoris.* We compared single nucleotide polymorphism (SNP) sharing and divergence across these samples by mapping all reads to the reference *C. rubella* genome, and incorporated genome resequencing data from 13 *C. grandiflora* accessions (25, 29) and 10 *C. orientalis* accessions (28) obtained previously. Because we are mapping our tetraploid sequences to a diploid, we expect to see high levels of apparent ‘heterozygosity’; as *C. bursa-pastoris* is highly selfing we expect the vast majority of these heterozygous sites to represent homozygous polymorphic differences between the two homeologous copies. Across the genome, both higher-order heterozygosity and depth appear to be quite uniform (Fig. S2A), suggesting the general absence of large-scale deletions and/or recombination events between homeologous chromosomes.

One category of particular interest is sites that exhibit ‘fixed heterozygosity’ across all of our *C. bursa-pastoris* samples, as these represent fixed differences between the subgenomes that arose either because of initial sequence differences between the parental species or due to subsequent mutations that have spread through the tetraploid species. We find that 58% of the 105,082 fixed heterozygous positions are fixed SNP differences between *C. orientalis* and *C. grandiflora.* Of the remainder, 30% represent sites that are segregating within *C. grandiflora,* only 0.2% are segregating within the highly selfing *C. orientalis,* while 0.06% are segregating in both species and 12% of fixed heterozygous sites are not found segregating in our diploid samples. Conversely, 95.4% of the 77,524 fixed differences identified between *C. orientalis* and *C. grandiflora* have both alleles in *C. bursa-pastoris* and 83% of these show fixed heterozygosity. The high proportion of fixed differences between diploids found within *C. bursa-pastoris* argues against widespread gene conversion having occurred between homeologous genes. However, there is spatial clustering of the 4.6% of sites fixed for only one of the alleles fixed between the diploids, suggesting a small degree of gene conversion between the subgenomes may have occurred (Fig. S2*B*). Finally, we examined patterns of transposable element insertion polymorphism sharing among the three species. As expected, principal component analysis places *C. bursa-pastoris* intermediately between the two diploid species (Fig. S3*A*), and 72% (2,415) of the insertions shared between two of the three species are shared between *C. bursa-pastoris* and *C. grandiflora,* 20% (675) are shared between *C. bursa-pastoris* and *C. orientalis*, while only 7% are shared uniquely between *C. grandiflora* and *C. orientalis*. These patterns are highly consistent with our earlier conclusions about hybrid origins, with the vast majority of fixed differences between subgenomes having been sampled either from the differences between *C. orientalis* and *C. grandiflora* or segregating variation from within the highly polymorphic outcrossing species.

To estimate the timing of origin of *C. bursa-pastoris,* SNPs from each individual were phased and identified as segregating on the *C. grandiflora* or *C. orientalis* descended subgenome (see Methods and SI text). Principal component analysis of these phased SNPs shows strong clustering of one subgenome (hereafter the ‘A subgenome’) with *C. grandiflora* and *C. rubella,* while the alternate subgenome (hereafter the ‘B subgenome’) clusters with *C. orientalis* (Fig. S3*B-C*). We used coalescent simulations applying the composite likelihood method implemented in fastsimcoal2.1 (30) to fit models of speciation to the joint site frequency spectra data from 60,225 phased SNPs at intergenic non-conserved regions and four-fold synonymous sites from the *C. bursa-pastoris* A subgenome, *C. grandiflora*, *C. bursa-pastoris* B subgenome and *C. orientalis.* We compared four different models using Akaike’s Information Criterion (AIC), including models with either a stepwise or exponential change in population size, and with or without post-polyploidization gene flow. We assumed a mutation rate of 7 * 10^−9^ per base pair per generation and a generation time of one year when converting estimates to units of years and individuals. The models show general agreement in terms of timing of origin, but the best-fit model is one without migration but with exponential growth following speciation (Fig. 2, see SI text). These analyses suggest that *C. bursa-pastoris* formed very recently, between 100,000 and 300,000 years ago, while the divergence time between the parental lineages of *C. orientalis* and *C. grandiflora* was considerably older, at approximately 900,000 years ago. Based on our simulations incorporating exponential population size changes we estimate that the current effective population size *(N_e_)* of *C. bursa-pastoris* (approx. 37,000-75,000) is considerably larger than that of the highly selfing, nearly invariant diploid *C. orientalis* (4,000), but much smaller than the *N_e_* of the highly outcrossing *C. grandiflora* (840,000). Given the shared variation between *C. bursa-pastoris* and both parental species, we can clearly rule out a single polyploid origin from two haploid gametes, although there is likely to be a high level of uncertainty about the precise number of founding lineages (20). Our best-fitting model implies that the *N_e_* of the founding population may be as large as 40,000. The reductions in *Ne* in *C. bursa-pastoris* and *C. orientalis* likely reflect the combined effects of demography and selfing, including the action of greater background selection in highly selfing populations.

**Figure 2.**
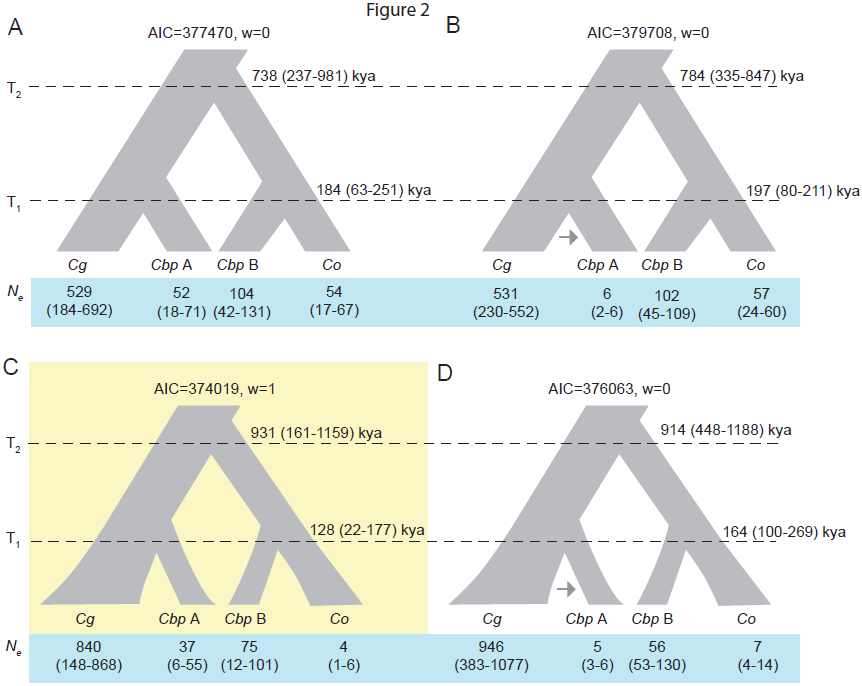
Demographic parameter estimates with 95% confidence intervals for four models of allopolyploid speciation in *Capsella.* Four models were investigated: A) Stepwise population size change, no gene flow, B) Stepwise population size change, asymmetric gene flow, C) Exponential population size change, no gene flow and D) Exponential population size change, asymmetric gene flow. Model C was preferred based on AIC and Akaike’s weight of evidence (w). Estimates of effective population sizes (*N_e_*) for *C. grandiflora (Cg), C. orientalis (Co), C. bursa-pastoris* (*Cb_p_* A and *Cb_p_* B) are given in thousands of individuals, and estimates of the timing of the origin of *C. bursa-pastoris* (*T_1_(Cbp)*) and the split between *C. grandiflora* and *C. orientalis (*T_1_(Cg-Co)*)* are given in kya. Note that for models with exponential population size change, *N_e_* corresponds to current *N_e_*. Confidence intervals are given in parentheses.

### Identification and Characterization of Putatively Deleterious Mutations

To examine evidence for gene loss, we looked for genes with significant reductions in normalized coverage in our 10 *C. bursa-pastoris* genomes compared with the diploid parental species, *C. grandiflora* and *C. orientalis.* We find only 76 genes with statistical evidence of gene loss specific to *C. bursa-pastoris*, suggesting that massive gene loss events have not yet occurred over approximately 200,000 years of divergence (Database S1). The identified deletions averaged 5,015 bp and spanned 1,067 to 19,643 bp in length, consistent with conclusions from ancient polyploids that most gene losses are mediated by short deletion events (31). While some of these deletions are in annotated genes that often show presence/absence polymorphism even in diploids, such as F-box proteins and disease resistance genes (32), others appear to be in essential genes (Database S1).

Since the coverage analysis requires significant reductions in read number across our entire set of *C. bursa-pastoris* individuals from a broad geographic range including Europe and Asia, it is likely to be highly enriched for deletions that are at high frequency or fixed across the species. To examine the potential for sample-specific deletions, structural variants were called using Pindel (33), and we identified deletions spanning 80% or more of a gene. In *C. bursa-pastoris*, 150 gene deletions unique to the species were identified, while 155 and 26 were found in *C. grandiflora* and *C. orientalis* respectively. The relative number of gene deletions compared to all other polymorphic deletions was moderately higher in *C. bursa-pastoris* than in the diploids (Chi-squared *p* < 0.05).

Given the apparently low levels of whole gene loss in *C. bursa-pastoris,* we were interested in examining whether more subtle signs of gene degeneration were apparent. We therefore assessed the numbers and frequencies of derived genic loss-of-function SNPs and insertion/deletion events (indels) causing frameshift mutations in coding regions. Possible compensatory mutations, including indels restoring frame and neighbouring SNPs with opposite effects (34) were also accounted for and removed from this analysis (see SI text).

We identified a large number of putatively deleterious SNPs in the *C. bursa-pastoris* genome (Table 1). Interestingly, a large proportion of these are fixed between subgenomes, implying that one of the duplicate copies may be inactive in all individuals. Nevertheless, we observe an excess of rare variants relative to 4-fold degenerate sites among the putatively inactivating mutations still segregating in *C. bursa-pastoris*, suggesting the continued action of weak purifying selection on at least some of these mutations (Fig. S4). To understand the origins of the putatively deleterious mutations, we examined the frequency and abundance of putatively inactivating mutations occurring in the diploid *Capsella* genomes (Table 1). Strikingly, the majority of fixed deleterious SNPs between the subgenomes in *C. bursa-pastoris* were also found in the diploid species; in particular, depending on the effect type, 40−60% of these were found in C. *orientalis,* 5-17% are in *C. grandiflora,* and 10-20% were found in both species.

**Table 1.** Counts of the different categories of SNPs segregating in *C. bursa-pastoris*^1^

| SNP effect | Unique to<br>*C. bursa-pastoris* | Shared with<br>*C. grandiflora* | Shared with<br>*C. orientalis* | Shared with<br>both |
| --- | --- | --- | --- | --- |
| Four-fold<br>synonymous | 10767 (1071) | 9203 (6875) | 9325 (5027) | 4106 (2755) |
| Start codon lost | 347 (22) | 75 (11) | 199 (71) | 125 (21) |
| Stop codon gained | 2829 (105) | 650 (55) | 1093 (348) | 485 (122) |
| Stop codon lost | 312 (18) | 138 (25) | 252 (105) | 220 (37) |
| Splice site lost | 1652 (74) | 394 (56) | 743 (271) | 454 (77) |

1- values in parentheses indicate fixed heterozygous SNPs likely reflecting fixed differences between homeologs

Nearly all (98%) of the putatively deleterious SNPs in *C. orientalis* also found in *C. bursa-pastoris,* are fixed in *C. orientalis,* reflecting its very low levels of polymorphism. Since *C. orientalis* is highly selfing and has a low *N_e_*, this may reflect the fixation of slightly deleterious mutations that have been passed on to *C. bursa-pastoris*. In strong contrast, many of the putatively deleterious SNPs from *C. grandiflora* are found at low frequencies in this highly polymorphic outcrosser, and less than 1% are fixed in *C. grandiflora.* Furthermore, there is evidence that the frequencies of these mutations have increased in *C. bursa-pastoris* compared to *C. grandiflora.* Thus, although there is an apparent bias towards *C. orientalis* being a source of deleterious SNPs, it is likely that additional ‘unique’ deleterious SNPs in *C. bursa-pastoris* originated from rare SNPs from the *C. grandiflora-like* ancestor. This means we may be underestimating the proportion of deleterious SNPs inherited from *C. grandiflora,* and overestimating the number of unique deleterious SNPs in *C. bursa-pastoris.* In either case, it appears that the beginnings of gene inactivation in *C. bursa-pastoris* may have had a ‘head start’ from both fixed slightly deleterious inactivations in *C. orientalis* and low-frequency, possibly recessive deleterious variation from the outcrossing *C. grandiflora*-like progenitor.

### Biased gene loss

To investigate bias in the types of genes being inactivated in all three *Capsella* species, we examined the functional categories of *Arabidopsis* orthologs using the Virtual Plant online server (35). Consistent with analyses of convergent ancient gene loss events (1), there was an enrichment of *C. bursa-pastoris* SNPs affecting stop codon gain and splice site loss in genes related to several functions associated with DNA (Table S1). Interestingly, segregating *C. grandiflora* stop codon gained SNPs are enriched for similar functional categories related to DNA repair and metabolism. These shared enrichments of functional categories are for mutations unique to each species as well as those shared between *Capsella* species, suggesting the similarities are not trivially due to shared polymorphisms. In contrast, putative loss-of-function mutations in *C. orientalis* were not enriched for any GO category. Thus, genes for which inactivating mutations segregate in the outbred diploid ancestor also tend to show independent, derived mutations causing gene inactivation in the recently formed tetraploid, but not in the highly homozygous selfing *C. orientalis*. However, genes within these enriched GO categories also tend to have larger coding regions on average (mean difference of 416 base pairs; Wilcoxon rank sum test *p* < 2.2 × 10^−12^) and as a result harbor more mutations in general. Since several of these GO categories have also been shown to be overrepresented for gene losses in ancient polyploids (1), mutational target size could be an underappreciated deciding factor in which genes are recurrently and rapidly lost following WGD. It is also possible that mutations in these genes are tolerated both in the outcrossing diploid and the allotetraploid, perhaps due to their predominantly recessive effects. This is consistent with the hypothesis that dosage–insensitive genes are more likely to experience gene loss following WGD (36), as these should also be more tolerated as polymorphisms in outbred diploid populations.

To test whether positive selection is driving the fixation of loss-of-function point mutations we scanned for signs of ‘selective sweeps’ or dips in neutral diversity surrounding fixed inactivating mutations. Average nucleotide diversity at 4-fold synonymous sites was assessed in 50kb windows of coding regions up- and down-stream of fixed putatively deleterious point mutations, separately for homeologous chromosomes. These windows show a large amount of variation, but no clear trend of reduced neutral diversity near focal mutations in comparison with synonymous substitutions (Fig. S5). Thus, there is no clear evidence that inactivating mutations were fixed by positive selection.

### No Evidence for Homeolog-Specific Loss of Expression

Genomic patterns suggest that there has not been rapid gene loss, and that the early stages of gene degeneration in *C. bursa-pastoris* may be driven in large part by ancestral variation from its diploid progenitors. However it is still possible that *de novo* changes in gene expression associated with epigenetic modification are widespread. To investigate this, we examined adult leaf gene expression patterns for the two homeologous genomes of one *C. bursa-pastoris* strain from our population genomic analyses and compared expression with replicate genotypes of the diploid progenitors (see SI text). In general, expression levels were comparable on the two subgenomes, with the median relative expression of 0.5 (Fig. 3A), showing no evidence of general expression biases towards one of the two homeologous genomes. We investigated if ancestral differences, or “legacies”, in gene expression could have contributed to differences in subgenome expression (see 14, 37). Overall there was conservation in levels of leaf gene expression between the diploids, but out of the 8164 genes assayed we identified 241 as differentially expressed (FDR adjusted *p* < 0.01). Genes overexpressed by at least two-fold in either diploid are also more highly expressed on the corresponding descendent subgenome (Fig. 3B), confirming that ancestral differences in gene expression predict differential expression in *C. bursa-pastoris*. Thus, as with our genomic analyses, we conclude that the early stages of homeologous gene silencing are determined at least in part by the gene expression history of diploid progenitors.

**Figure 3.**
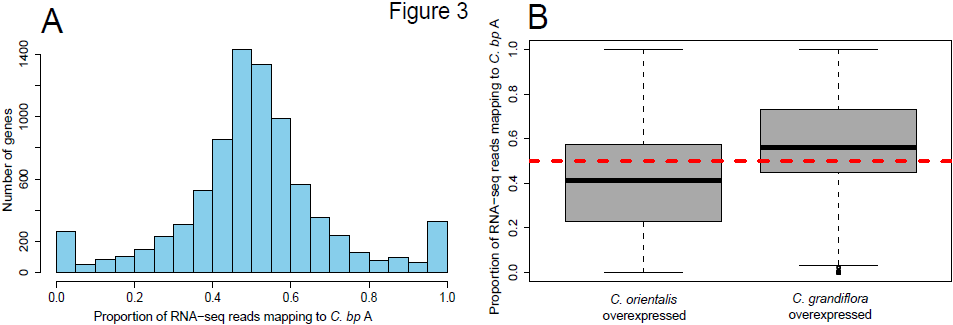
Transcriptome analyses testing for homeolog-specific expression. A) Proportion of reads mapping to *C. bursa-pastoris* subgenome A for all genes assayed. B) Proportion of reads mapping to *C. bursa-pastoris* subgenome A in genes overexpressed within each diploid progenitor. The dashed red line indicates where an equal proportion of reads maps to both *C. bursa-pastoris* subgenomes.

### Quantifying Relaxation of Purifying Selection

Overall, we detected small numbers of gene loss events, and signs that many major effect mutations were inherited from parental species. This suggests that the earliest stages of polyploid gene degeneration may be subtle, and reflect a global shift in selection pressures genome-wide rather than rampant gene loss. To investigate the strength of selection acting on *C. bursa-pastoris* compared to its close diploid relatives, we estimated the distribution of fitness effects (DFE) of deleterious mutations (38) for all three *Capsella* species using allele frequency spectra and separating the two homeologous genomes of *C. bursa-pastoris.* The method explicitly incorporates non-equilibrium population size changes, and thus should account for between-species differences in demographic history. We estimated the DFE at both 0-fold non-synonymous sites and conserved noncoding sites as previously defined through conservation between the *Capsella* reference and nine additional Brassicaceae species (29). Between *C. bursa-pastoris* subgenomes there was no significant difference in the DFE for either site type, providing no evidence for early signs of biased fractionation in terms of the efficacy of selection on the two subgenomes (Fig. 4). However, genes showing homeolog-specific gene silencing (see SI text) are under significantly less purifying selection on the silenced subgenome *(p* < 0.05), suggesting that the ancestral differences in gene expression inherited in *C. bursa-pastoris* can help drive the early degeneration of lower-expressed homeologous genes.

**Figure 4.**
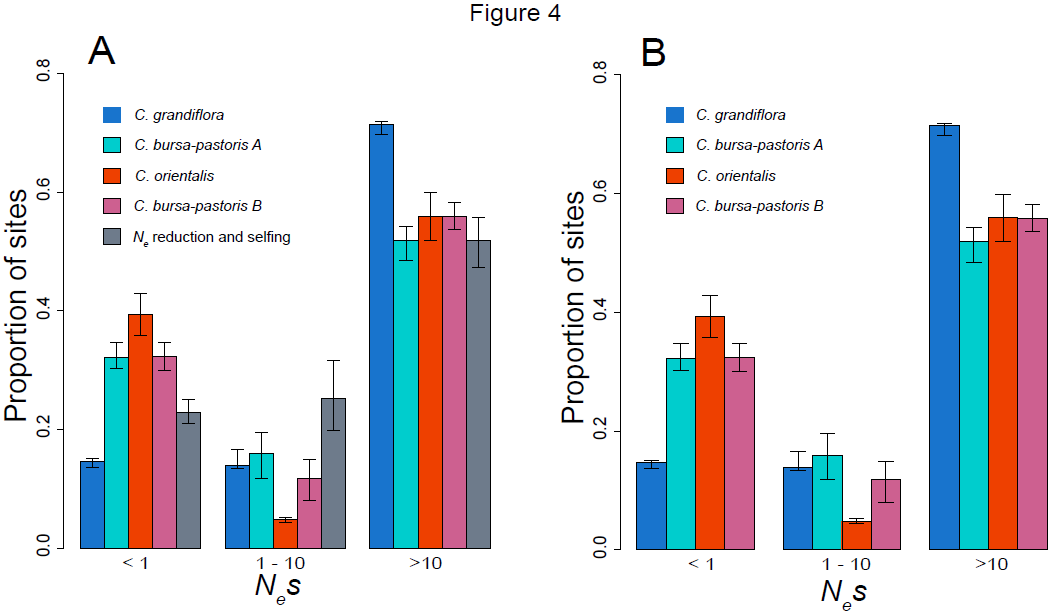
Estimates of the distribution of fitness effects (DFE) of deleterious mutations at 0-fold nonsynonymous sites (A) and conserved noncoding sites (B), based upon the allele frequency spectra of SNPs at these sites. The simulated DFE for 0-fold non-synonymous sites under effective population size (*N_e_*) reduction and a shift to selfing is also shown in A. The strength of selection is measured as *N*_*e*_*s*, where s is the strength of selection. Error bars correspond to 95% confidence intervals of 200 bootstrap replicates of 10 kilobase blocks.

As predicted, the *C. bursa-pastoris* A subgenome showed a significantly elevated proportion of effectively neutral mutations (*N*_*e*_*s* < 1, where *s* is the strength of selection against deleterious mutations) compared with *C. grandiflora* for both nonsynonymous and conserved noncoding sites, consistent with the prediction that the transition to polyploidy has led to genome-wide relaxed selection (Fig. 4A. In contrast, however, *C. orientalis* shows a significantly elevated proportion of effectively neutral nonsynonymous mutations compared with the *C. bursa-pastoris* B subgenome (Fig. 4B, and no difference between the two for conserved noncoding sites (Fig. 4B. Given the severe reduction in *N*_*e*_ and selfing mating system in *C. orientalis* (Fig. 2), this implies that demographic effects and mating system changes could have similar effects on the strength of purifying selection as the ploidy transition.

To what extent is relaxed selection in *C. bursa-pastoris* driven by the shift to self-fertilization and reductions in *N_e_*, rather than the change in ploidy? To address this question, we conducted forward simulations, modeling a diploid population that has experienced the mating system transition and reduction in *N_e_* over the timescale inferred for *C. bursa-pastoris* (Fig. 2; see SI Text). In these simulated populations, we observe a considerable increase in the proportion of effectively neutral mutations (Fig. 4A, implying that the shift in the strength of purifying selection is due in part to greater effects of drift caused by demography and selfing.

However, the extent of relaxed selection due to the reduced *N*_*e*_ alone is considerably smaller than the one we observe (Fig. 4); based on the bootstrapped 95% CI, the contributions of demography and changes in mating system account for 30-66% of the shift in purifying selection in *C. bursa-pastoris*. Thus, this implies that the transition to polyploidy and the corresponding redundancy in gene function is also contributing to the changes in selection.

### Conclusions

Our results provide comprehensive evidence for recent allopolyploid origins of *Capsella bursa-pastoris* from two parental lineages ancestral to present-day *C. orientalis* and *C. grandiflora.* Given the patterns of polymorphism and expression, our results suggest that gene degradation and silencing following allopolyploidization can get a head start from standing variation in progenitors. By contrasting patterns of functional polymorphism and divergence genome-wide, we find evidence for significant relaxation of purifying selection driven by both increased drift and relaxed selection due to gene redundancy, but no evidence for massive gene loss rapidly upon polyploidization. Demographic and historical factors that accompanied polyploidy were responsible for a large proportion of the inferred relaxation of purifying selection, suggesting that masking due to gene redundancy is only one of several contributors to early genome evolution following polyploidy. Taken together, and in contrast with patterns inferred in other systems (9, 11) our results suggest that the early stages of allopolyploid evolution in *C. bursa-pastoris* were characterized by relaxed purifying selection rather than large-scale "genomic shock" and rapid gene loss.

### Materials and Methods

Seeds from ten *Capsella bursa-pastoris* plants were collected from one individual per population across the species range in Eurasia (see SI text). Nuclear genomic DNA was extracted from leaf material for the 10 *C. bursa-pastoris* individuals using a modified CTAB protocol (39). Paired-end 108 bp and 150 bp reads were generated for *C. bursa-pastoris,* with median depth coverage of 20 reads per site using Illumina GAII and Illumina HiSeq 2000 (Illumina, San Diego CA, USA). Sanger sequencing of a more extensive sample of *C. bursa-pastoris* was used in combination with published sequences from other *Capsella* species (see SI text) to validate the conclusions about hybrid origins. For the transcriptome analyses, we extracted RNA from adult leaf tissue with a Qiagen Plant RNEasy Plant Mini Kit from one *C. bursa-pastoris* individual (SE14), six *C. grandiflora* individuals, and three biological replicates each of two *C. orientalis* individuals. RNA sequencing libraries were generated using the TruSeq RNA v2 protocol and sequenced on Illumina HiSeq 2000.

*De novo* fragment assemblies for *C. bursa-pastoris* were generated using Ray v 1.4 (40). Read mapping of Illumina genomic reads to the *C. rubella* reference genome (21) was conducted using Stampy version 13 (41), and phasing of read-mapped samples was conducted using HapCUT (42). SNP calling and polymorphism analyses were conducted using the Genome Analysis Toolkit (43). We constructed phylogenies using RAxML’s (44) rapid bootstrap algorithm to find the best-scoring ML tree. For demographic inferences, we used fastsimcoal2.1 (30) to infer demographic parameters based on the multidimensional site frequency spectrum for *C. grandiflora*, *C. orientalis*, and the two *C. bursa-pastoris* homeologous genomes. To evaluate the fit of the best demographic model (Table S2), simulated datasets were compared with the observed site frequency spectra (Fig. S6). The DFE was estimated for each species using the maximum likelihood approach designed by Keightley and Eyre-Walker (38). We conducted forward simulations using the software SLiM (45). SnpEff (46) was used to predict the effects of SNPs called using GATK. See SI Text for detailed methods on *de novo* assembly, expression analysis, population genomics analyses and validation.

## ACKNOWLEDGEMENTS

This project was funded by an NSERC Discovery Grant and a Genome Quebec/Genome Canada grant to S.I.W., and grants from the Swedish Research Council to T.S. and to M.L. G.M.D. was supported by an NSERC scholarship, and E.B.J. by an NSF graduate fellowship. Fastsimcoal2.1 computations were performed on resources provided by SNIC through Uppsala Multidisciplinary Center for Advanced Computational Science (UPPMAX) under Project b2012190. DNA and RNA sequencing was performed by the Genome Quebec Innovation Centre, and RNA sequencing by the SNP&SEQ Technology Platform, Science for Life Laboratory at Uppsala University, a national infrastructure supported by the Swedish Research Council (VR-RFI) and the Knut and Alice Wallenberg Foundation. We thank Vitor Sousa at University of Bern for help with Fastsimcoal analyses.

